# Host-pathogen-vector continuum in a changing landscape: Drivers of *Bartonella* prevalence and evidence of historic spillover in a multi-host community

**DOI:** 10.1101/2023.09.25.559241

**Authors:** B.R. Ansil, Ashwin Viswanathan, Vivek Ramachandran, H.M. Yeshwanth, Avirup Sanyal, Uma Ramakrishnan

**Affiliations:** National Centre for Biological Sciences, Tata Institute of Fundamental Research, Bangalore, Karnataka, India, 560065; Manipal Academy of Higher Education, Manipal, Karnataka, India, 576104; Nature Conservation Foundation, Mysore, Karnataka, India, 570017; Trivedi School of Biosciences, Ashoka University, Sonipat, Haryana, 131029

**Keywords:** Spillover, Western Ghats, Small mammals, Vector-borne pathogens

## Abstract

Small mammals and their ectoparasites present a unique system to investigate the eco-epidemiology of multi-host vector-borne pathogens and identify specific bacterial spillover determinants. We applied ecological and evolutionary analyses in a rainforest-human-use mosaic to investigate *Bartonella spp.* across small mammal and ectoparasite communities. We observed substantial overlap among small mammal communities in different habitat types, predominantly driven by habitat generalists. Most ectoparasites were generalists, infecting multiple hosts. We observed high *Bartonella* prevalence at both study sites –a forest-plantation mosaic (47.4%) and a protected area (28.8%). Seven of the ten ectoparasite species sampled were also positive for *Bartonella*, following prevalence trends in their hosts. A generalised linear model revealed an independent association between aggregated ectoparasite load in hosts and *Bartonella* prevalence, implicating ectoparasites in transmission. *Bartonella* lineages from small mammals were host-specific, while ectoparasites carried *Bartonella* associated with other small mammal hosts, indicating the potential for cross-species transmission. Phylogenetic ancestral trait reconstruction of *Bartonella* haplotypes suggest historic spillover events in the small mammal community, validating the potential for contemporary spillover events. These results highlight the necessity to disentangle the complex relationship between hosts, ectoparasites, and pathogens to understand the zoonotic implications of undetected spillover events in such multi-host communities.

## 1. Background

Large-scale modification of natural habitats is associated with loss of biodiversity and ecosystem services [1]. Other unexpected consequences, such as an elevated risk of zoonotic infections, are poorly acknowledged and understood, especially in biodiverse tropical systems. Both theoretical and empirical studies demonstrate that land-use change and associated habitat perturbations can result in modified communities of zoonotic hosts driven by movement of animals across habitat boundaries [2,3].Consequently, there is potential for pathogen spillover between animals that did not previously co-occur, i.e. from reservoir hosts to incidental hosts, directly or indirectly through intermediate hosts and vectors [4,5].

Spillover between wildlife can lead to the emergence of zoonoses [4], and has been implicated in several past epidemics, including SARS-CoV-1 [6]. Pathogen sharing between animal host communities can lead to the recombination and evolution of new variants with altered host ranges, vector specificities, and pathogenic potential [7]. Spillover events, however, are complex and infrequent, influenced by numerous ecological and evolutionary filters that vary in space and time, including the distribution of reservoir hosts, pathogen prevalence, and infection intensity [8]. For example, a high density of reservoir hosts elevates direct interactions, leading to density-dependent transmission among the reservoir population and to other susceptible members of the community [9]. Additionally, in the case of vector-borne pathogens, ectoparasites also respond to host densities and influence pathogen prevalence in reservoirs and incidental hosts [10].

Bats and small mammals (rodents and shrews) are the best-known reservoir taxa for zoonotic pathogens [11]. Small mammals, however, harbour a high number of pathogens [12,13] owing to their disproportionately high diversity within mammalian communities [14], even more so than bats. Small mammals also differ from bats in that they often increase (rather than decrease) in abundance in altered landscapes, resulting in the elevated prevalence and transmission risk of associated zoonoses [15,16]. Small mammal densities are known to increase locally, for example, when large herbivore densities decrease [17], leading to an increase in pathogen prevalence [18]. In addition, small mammals with generalist natural histories (often reservoirs) tend to move across habitat boundaries where they interact with incidental hosts [4,19], thereby moving pathogens across these boundaries. We expect such processes to be especially prevalent in tropical regions, as they harbour some of the richest small mammal communities [20], the highest rates of habitat modification [21,22], and therefore the highest predicted risk of zoonotic infections [23,24].

To understand the zoonotic consequence of small mammals and their ectoparasites in changing landscapes, we must first understand how their communities are structured and how they have responded (and continue to respond) to land-use change and habitat modification. In this study, we characterize differences in host community structure, ectoparasite community structure, and pathogen prevalence in a human-dominated rainforest-plantation landscape in South India, part of the Western Ghats biodiversity hotspot. We used *Bartonella*, a generalist zoonotic bacterial pathogen, as a model system to understand variation in prevalence and transmission dynamics in the landscape. We sampled small mammal communities and their ectoparasites in this landscape to address the following questions – i) How are small mammal communities structured in a tropical rainforest-plantation landscape? ii) How does *Bartonella* prevalence vary across small mammal species in this human-use landscape? iii) What are the main drivers of *Bartonella* prevalence and transmission in the landscape? (iv) and finally, do we find evidence for spillover, detected as shared *Bartonella* haplotypes between members of these communities? We began this study in a forest-plantation mosaic and expanded to a relatively less disturbed (currently) protected area. However, the protected area had significant human-use in the past [25], similar to the forest-plantation mosaic, allowing us to derive insights that can be broadly applied and generalized. Given the mixed-use nature of the landscape, we expected similarities between small mammals and ectoparasite communities across land-use types [26]. We hypothesized that *Bartonella* prevalence would be high and significantly influenced by the densities of known host species (e.g., *Rattus satarae*) [9,27]. We also expected that *Bartonella* haplotypes would be specific to host species [28] with limited spillover (vector-mediated) among hosts that occupy the same habitats [29].

## 2. Methods

### (a) Field sampling

We carried out this study in two sites in the central Western Ghats: i) Kadamane, a mixed-use, forest-plantation mosaic in the Karnataka state in India, and ii) Kudremukh National Park, a protected area in the same state (**Figure 1**). We sampled small mammals (rodents and shrews) from different land-use types in Kadamane and Kudremukh between 2016 and 2021. Kadamane was sampled three times during this period (2016, 2017, and 2018) and Kudremukh was sampled once in 2021. In Kadamane, we sampled in forest fragments, grasslands, tea plantations, rested plantations (unmanaged cardamon and coffee), and built-up (human habitation). In Kudremukh, we sampled forests, grasslands, and built-up (a partially occupied township inside Kudremukh National Park). In both sites, sampling was conducted during dry months between January-April.

**Figure 1:**
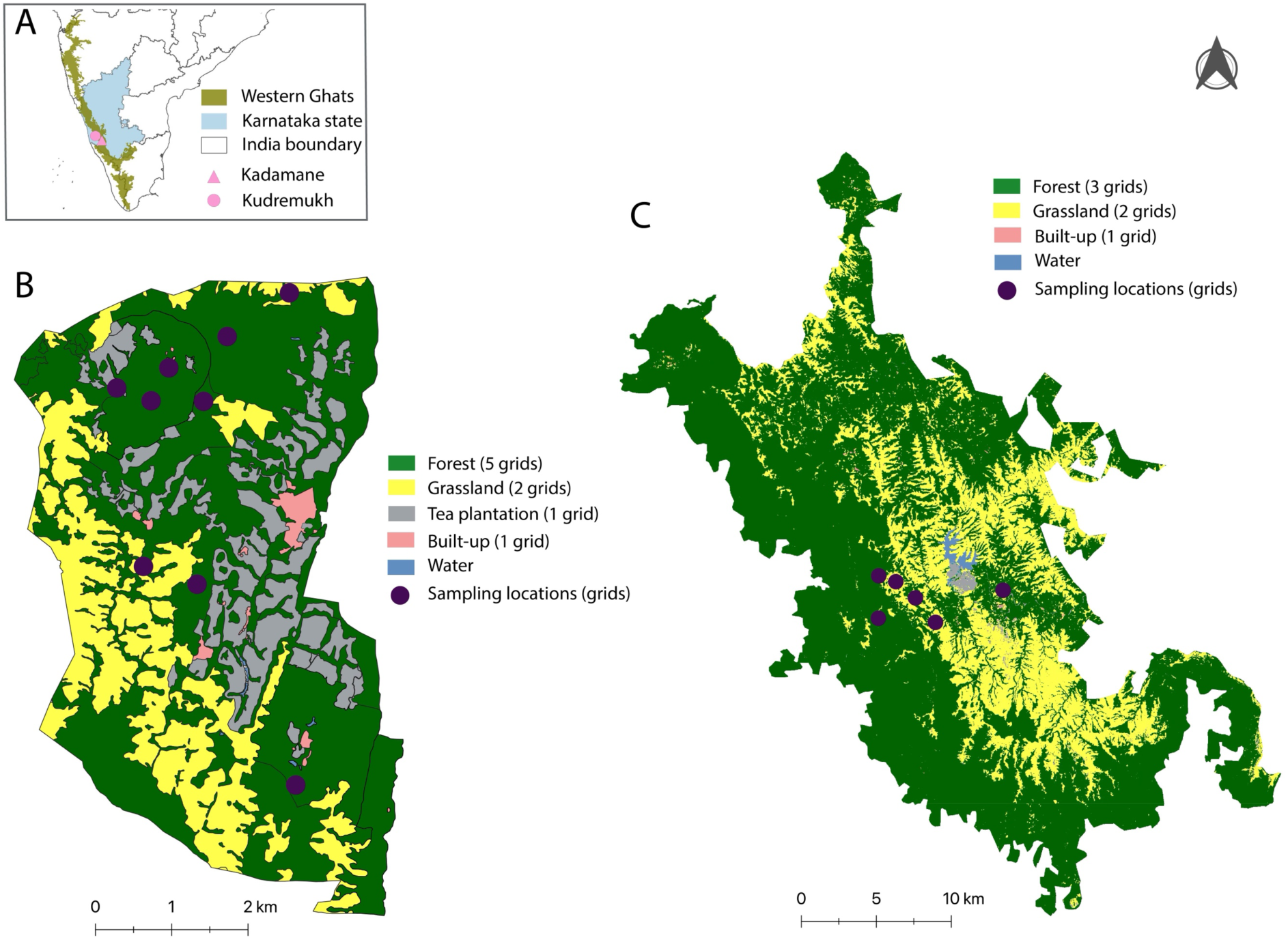
Map of the study area. A) Map of India with Karnataka state, Western Ghats and study sites marked. (B) Land-use map of Kadamane forest-plantation mosaic with sampling locations marked. (C) Land-use map of Kudremukh National Park with sampling locations marked. The legends indicate different land-use types and the number of grids sampled per land-use type. Note that rested plantations are grouped under forests.

We used medium-sized Sherman traps in a grid framework to sample small mammals. A trapping grid consisted of 10 trapping lines, each line consisting of 10 traps separated by 10 meters, covering an area of one hectare in total in each habitat type. An additional 20 traps were placed on trees to capture arboreal species in forests and rested plantations. Trapping efforts were proportionately divided based on the area available under each land-use category. Specifically, in Kadamane, this translated to three grids in forests, two grids in grasslands, two grids in rested plantations, one grid in tea plantations, and one grid in built-up areas. Because the rested plantations in Kadamane were often indistinguishable from forests, we grouped them under ’forests’ in all analyses. In Kudremukh, three grids were placed in forests, two in grasslands, and one in built-up areas (**Figure 1**). Each grid was sampled for four consecutive days during each trapping session (assuming a closed population). All traps were baited using peanut butter and locally available seeds. Traps were checked twice a day (morning and evening) for captured individuals. Captured individuals were anesthetised using Isoflurane for collecting measurements and samples. Small mammal species were first identified in field with the help of taxonomic keys and field guides [30–32]. In addition, ear clippings were collected to confirm species identification through molecular methods. We collected dried blood spots from every captured individual in 2018 and 2021 to screen for pathogens. To detect and collect ectoparasites, captured individuals were combed on a white paper for 2-3 minutes. All individuals were marked and released at the site of capture after sample collection (except in 2018; individuals were sacrificed for harvesting organs to screen for various pathogens).

### (b) Small mammal community

DNA was extracted from ear clippings using DNeasy Blood & Tissue Kit (Qiagen), following which 1140 bp of Cytochrome b (*Cytb*) gene was amplified and sequenced [33]. Maximum likelihood phylogenetic trees were constructed to identify cryptic *Mus* species and validate morphological species identification. All the other species were identified morphologically and validated using BLAST (blast.ncbi.nlm.nih.gov) similarity scores of *Cytb* sequences.

Based on species identification, we created a species abundance matrix for each land-use type, followed by a distance matrix using the Morisita-Horn index, a standard beta diversity measure to calculate dissimilarity [34]. The beta diversity index was plotted following hierarchical clustering to visualise the dissimilarity between small mammal communities in different land-use types and sampling grids. We also calculated the relative abundance of each small mammal species in each land-use type to understand species-habitat associations. The community ecology package Vegan [35] in R was used for this analysis.

We used a Capture-Mark-Recapture approach [36] to estimate densities of the four most common small mammals by creating individual capture histories in forest and grassland. We assumed a closed population with varying capture probability to estimate population size following a customised log-linear model [37]. Density (number of individuals per hectare) for each species was estimated after accounting for the number of grids sampled in each habitat. We used R package *Rcapture* for this analysis [37].

### (c) Ectoparasite community

Ectoparasites were brought back to the laboratory in 100% ethanol. Mites, ticks, and fleas were segregated using a stereo-zoom microscope. We identified ticks and fleas to the genus level under a high-resolution microscope with the help of taxonomic keys and monographs [38–40]. Mites were mounted on a slide for observation under the microscope and identified through descriptions in a monograph [41]. We serially named and counted individual ectoparasites that belonged to the same morphotypes (potentially species). We constructed a distance matrix based on ectoparasite occurrence per individual small mammal species [42]. We used this matrix for a Principal Coordinate Analysis (PCoA) in R [43], and plotted the clustering using ggplot2 [44] to understand the overlap in ectoparasite communities between small mammal species (the first two PCoA axes). We also estimated ectoparasite density, or the mean number of each morphotype per host species in forests, grasslands, and built-up areas. The infestation rate, i.e., frequency of ectoparasite (mites, ticks, and fleas) occurrence in each host species and habitat, was recorded as aggregated ectoparasite load.

### (d) *Bartonella* screening and prevalence estimation

To detect *Bartonella* in small mammals, DNA was extracted using QIAamp DNA Mini Kit (Qiagen) from the dried blood spots (DBS) collected in 2018 (Kadamane) and 2021 (Kudremukh), and screened for *Bartonella* specific *rpoB* gene (825 bp) using conventional PCR [45]. *Bartonella* prevalence in the Kadamane samples was reported previously [27]. To detect *Bartonella* in ectoparasites, ectoparasite were pooled, mechanically homogenised, and DNA was extracted using DNeasy Blood & Tissue Kit (Qiagen). A pool contained 1-10 individual ectoparasites (mean = 3.06) belonging to the same species (previously called morphotypes) collected from the same individual small mammal. We used PCR reagents as follows-25 μl reaction mixture composed of 6 µl of template DNA, 12.5 µl of 1X HotStarTaq Master Mix (Qiagen), 1 µl each of 5 μM primers and 4.5 µl of nuclease-free water. PCR products were visualized on a 1.5% agarose gel stained with GelRed Nucleic Acid Gel Stain (Biotium Inc) with reference to negative and positive controls set along the test samples. Positive samples were purified using AMPure XP magnetic beads (Beckman Coulter) and sequenced. Sequences were verified for *Bartonella* after comparing with the reference database (NCBI) using BLAST [46]. All *Bartonella* sequences generated in this study have been deposited in GenBank under accession numbers OR574780-OR574826.

Prevalence was calculated as the proportion of positive individuals against the number of individuals tested. While calculating the overall prevalence, we included two forest grids from Kudremukh twice (randomly selected) to compensate for additional trapping effort in Kadamane (rested plantations) to make the sites comparable. We calculated 95% confidence intervals (Wilson interval) associated with the prevalence using the R package *Hmisc* [47]. Prevalence (positivity) in ectoparasite pools was represented as the proportion of positive pools among the total number of ectoparasite pools tested.

### (e) Ecological correlates of *Bartonella* prevalence

We used a generalized linear model (GLM) to understand drivers of *Bartonella* prevalence among small mammals in the landscape. We modelled prevalence as the incidence of *Bartonella* (presence/absence) in each individual small mammal as a function of the sampling site, aggregated ectoparasite load (the proportion of all individuals of the species with any ectoparasite presence - mites, ticks, and fleas - within the relevant habitat-site combination), and the density of the small mammal species itself. We assumed a binomial error distribution as our response consisted of 0s and 1s. We restricted this analysis to the 10 species-habitat-site combinations that contained data from more than three individuals of the species, together comprising 196 individuals of five species. We also examined host species identity and habitat type as covariates but dropped them because the models did not converge. The covariate ’aggregated ectoparasite load’ is the frequency of ectoparasite occurrence in each species and habitat, and is not ectoparasite density per individual small mammal. We considered this a more reliable measure than raw ectoparasite densities since it is possible to occasionally encounter individuals with heavy infestation in the community and this might potentially influence our inference.

### (f) Phylogenetic analyses

We generated a consensus of forward and reverse sequences for the partial *rpoB* sequence. These sequences were aligned with known *Bartonella* species sequences from the NCBI database and sequences reported in Ansil et al 2021 (accession number MT787671-MT787733) [27]. A 773 bp alignment was created using MUSCLE in MEGA 7.0.26 [48]. Sequences generated from the ectoparasites were also included in this alignment. Alignment was codon-optimized and tested for the appropriate nucleotide substitution models using PartitionFinder 2.1.1 [49]. We performed a Bayesian phylogenetic analysis using this alignment in BEAST 1.8.4 [50]. The phylogeny was run for 10^8^ Markov Chain Monte Carlo (MCMC) cycles. Effective sample sizes (ESS) and parameter convergence of MCMC sampling were assessed using Tracer 1.6.0 [51]. The resulting trees were summarised into a consensus tree after 25% burnin, and were visualized and annotated using FigTree 1.4.4 [52]. The phylogenetic tree was rooted using *Brucella melitensis* (AY562181) as an outgroup. Internal nodes of the tree are marked and coloured based on the Posterior Probability (PP).

### (g) Host- *Bartonella* association

We also constructed a tanglegram (co-phylogeny) using sequence data from the hosts (Cytb) and *Bartonella* (*rpoB*) in TREEMAP [53] to understand host-*Bartonella* haplotype association. We visually analysed the resulting co-phylogeny for potential host switching (spillover) events.

A separate alignment was constructed using unique *Bartonella* haplotypes and considered for a Bayesian phylogenetic ancestral trait reconstruction. We restricted this analysis to unique haplotypes recovered from each host species. We included one additional sequence from *Mus cf. fernandoni* to avoid over-representation and subsequent biases from *Rattus satarae* sequences. We used six identified host states as traits to estimate host probabilities for all ancestral nodes in BEAST 1.8.4 [50]. The resultant phylogeny was summarised and annotated as described before.

## 3. Results

### (a) Small mammal community

We recorded 11 small mammal species, of which nine were rodents, and two were shrews **(Table S1).** The rodent species were *Rattus satarae*, *Rattus rattus*, *Mus cf. fernandoni*, *Mus cf. famulus*, *Mus cf. terricolor*, *Golunda ellioti*, *Vandeleuria nilagirica*, *Platacanthomys lasiurus*, and *Funambulus tristriatus*. The two shrew species were *Suncus niger* and *Crocidura horsfieldii*. The three *Mus* species were morphologically similar and could only be differentiated using their phylogenetic clustering **(Figure S1)**. The average genetic distances (p distances) for *M. cf. fernandoni*, *M. cf. famulus*, and *M. cf. terricolor* from their respective parent species (for example, *M. cf. fernandoni* vs *M. fernandoni*) were 0.05, 0.07 and 0.09, respectively.

Among the various habitats sampled, grasslands had the most diverse small mammal communities with five species detected in both Kadamane and Kudremukh. Forest fragments in Kadamane also had five species of small mammals, but forests in Kudremukh only had three species. Built-up areas were less diverse with just three species captured at each site, while tea plantations were the most species-poor land-use type, with only one species (*M. cf. famulus*) captured once in 2016.

Based on the hierarchical clustering of beta diversity, forests (FR16, FR17, FR18, and FR21) showed similar community assemblages by forming a single cluster with minimal variability across years (<20%); the forest community from Kudremukh (FR21) was also a part of this cluster (**Figure 2A** **left)**. The grassland and tea plantation (TP16) formed a cluster with 40% dissimilarity with the forest cluster. The grassland communities had minimal variability (<20%) between years (GL16, GL17, GL18) and sites (GL21). Built-up habitats were the most dissimilar (60%) to other land-use types, although they retained two species (*S. niger* and *M. cf. famulus*) detected in the other land-use types (**Figure 2A** **right)**. Overall, Overall, the small mammal community composition within each habitat was similar across years.

**Figure 2:**
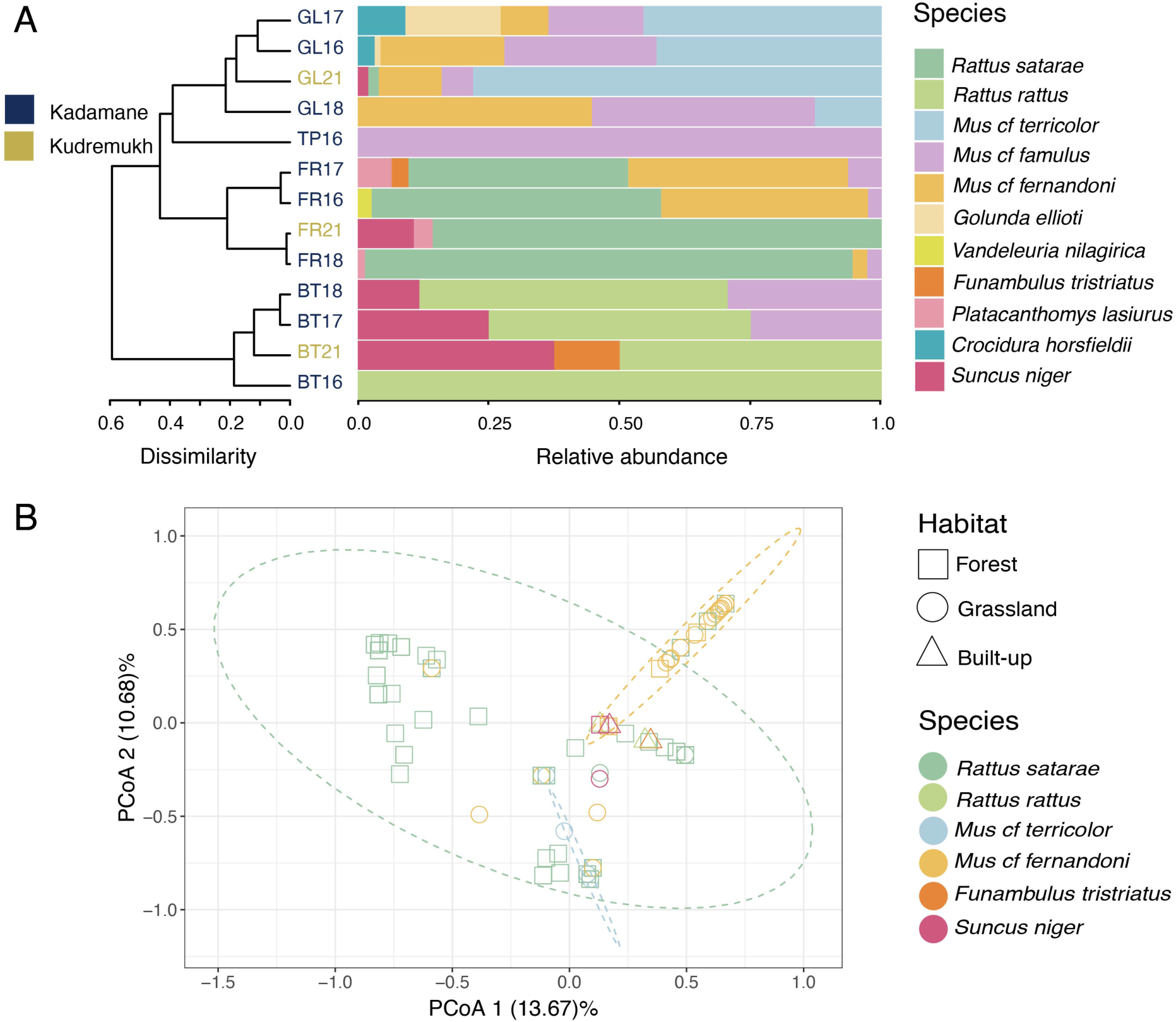
Community structure of small mammals and their ectoparasites. A) Left: a dendrogram showing dissimilarity between small mammal communities inhabiting different habitats sampled and sampling years (2016, 2017, 2018, 2021). Abbreviations: FR-Forest, GL-Grassland, TP-Tea plantations, and BT-Built-up areas. Right: Relative abundance of different small mammal species captured in different habitats. B) Principal Coordinate (PCoA) plot showing ectoparasite community structure across different small mammals and habitats. The ellipses represent the 95% confidence level of the respective sample group centroid.

We were able to estimate densities for four (most abundant of the eleven) common small mammal species (*Rattus satarae*, *M. cf. fernandoni*, *M. cf. famulus*, and *M. cf. terricolor*). These densities are reported in Supplementary results and **Figure S2**.

### (c) Ectoparasite community

Ten ectoparasite species were recorded from the small mammals; two species of mites belonging to the genus *Laelaps*, three species of ticks belonging to the genus *Rhipicephalus*, *Ixodes* and *Haemaphysalis* and five species of fleas belonging to the genus *Xenopsylla* **(Table S2)**. Since more than one species was detected under *Laelaps* (mites) and *Xenopsylla* (fleas), these morphotypes were named serially *(e.g. Xenopsylla sp 1, Xenopsylla sp 2* and so on). Only one morphotype each was identified from three tick genera. Interestingly, six ectoparasite species (60%) were detected in multiple hosts (generalists), and six of eight small mammals sampled (75%) were polyparasitic (infested with two or more ectoparasite species).

Our PCoA revealed substantial overlap and variation in ectoparasite communities hosted by small mammals (**Figure 2B**). *R. satarae* showed significant variation in ectoparasite community along the PCoA axis 1 and 2 (13.67 % and 10.68 % explained, respectively), while *M. cf.* fernandoni predominantly displayed limited variation (similar ectoparasite community dominated by *Laelaps* 1). However, individuals from *R. rattus*, *F. tristriatus*, *M. cf. fernandoni*, *M. cf. terricolor*, and *S. niger* showed ectoparasite community similarity with *R. satarae*. We also estimated the mean density of each ectoparasite species recovered from each small mammal species per habitat presented in Supplementary results and **Figure S2**.

### (d) Bartonella prevalence in small mammals

We detected *Bartonella* in five species of small mammals from both sites; *R. satarae*, *R. rattus*, *M. cf. fernandoni*, *M. cf. famulus,* and *S. niger* (**Table 1**) with varying prevalence (0-75.8%). In both sites, *R. satarae* showed the highest prevalence of 75.8% (n= 66) in Kadamane and 72% (n= 25) in Kudremukh, followed by *M. cf. fernandoni* that showed a low prevalence of 54.5% (n=22) in Kadamane and 40% (n=5) in Kudremukh. *M. cf. famulus*, on the other hand showed a low prevalence of 3.8% in Kadamane (n=26, only one individual positive) and zero prevalence in Kudremukh (n=3), while *M. cf. terricolor* showed zero prevalence at both sites despite good sample size (n =6 and 37 in Kadamane and Kudremukh, respectively). Other species, such as *R. rattus* and *S. niger* had low prevalence, but also had low sample sizes, hence we have minimal confidence in their prevalence estimates.

**Table 1:**
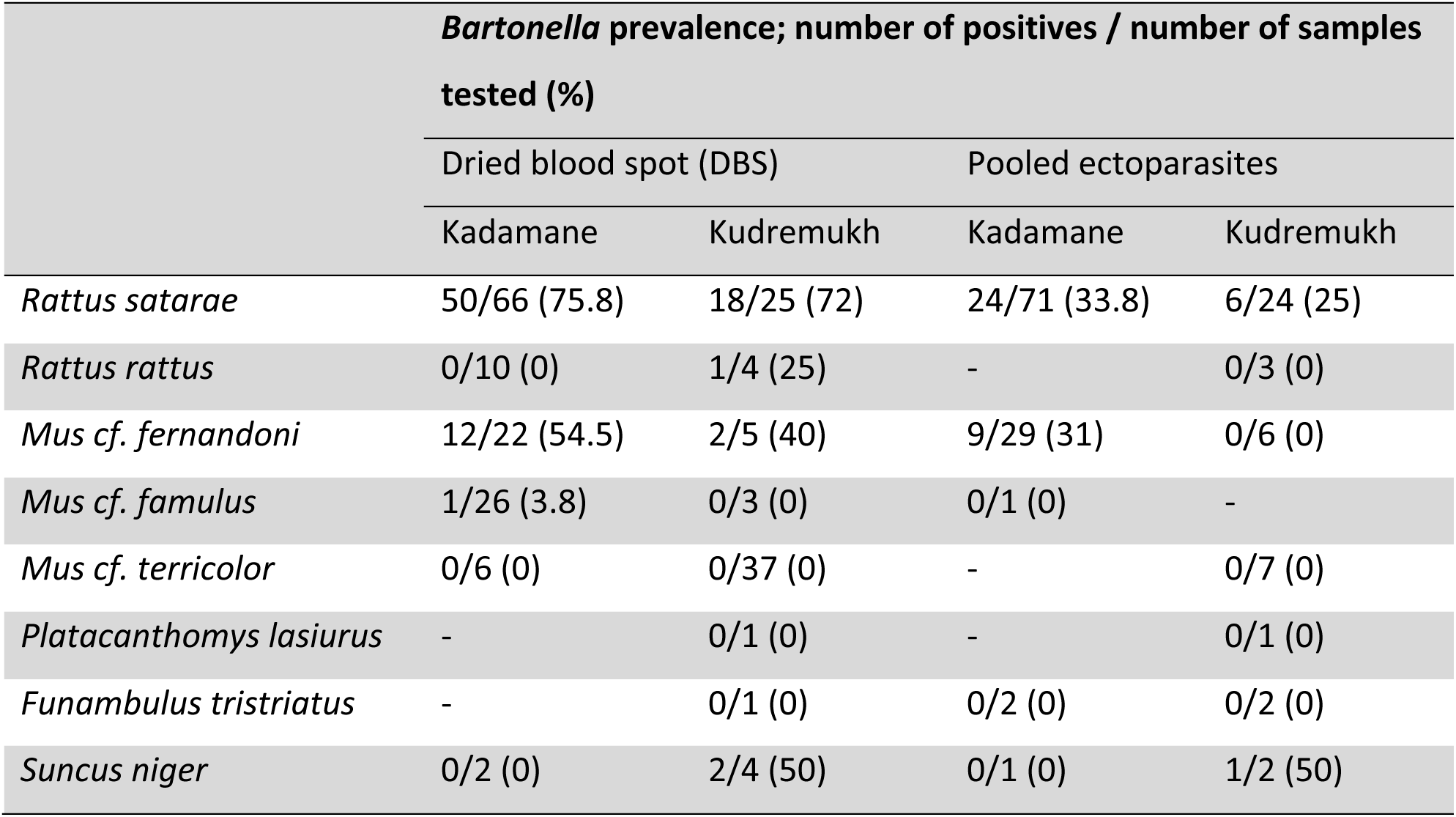
*Bartonella* prevalence in small mammal blood and ectoparasites. Prevalence reported for ectoparasites is not true prevalence since ectoparasites were pooled for screening (see methods).

|  | *Bartonella* prevalence; number of positives / number of samples tested (%) |  |  |  |
| --- | --- | --- | --- | --- |
|  | Dried blood spot (DBS) |  | Pooled ectoparasites |  |
|  | Kadamane | Kudremukh | Kadamane | Kudremukh |
| *Rattus satarae* | 50/66 (75.8) | 18/25 (72) | 24/71 (33.8) | 6/24 (25) |
| *Rattus rattus* | 0/10 (0) | 1/4 (25) | - | 0/3 (0) |
| *Mus cf. fernandoni* | 12/22 (54.5) | 2/5 (40) | 9/29 (31) | 0/6 (0) |
| *Mus cf. famulus* | 1/26 (3.8) | 0/3 (0) | 0/1 (0) | - |
| *Mus cf. terricolor* | 0/6 (0) | 0/37 (0) | - | 0/7 (0) |
| *Platacanthomys lasiurus* | - | 0/1 (0) | - | 0/1 (0) |
| *Funambulus tristriatus* | - | 0/1 (0) | 0/2 (0) | 0/2 (0) |
| *Suncus niger* | 0/2 (0) | 2/4 (50) | 0/1 (0) | 1/2 (50) |

We observed similar patterns of *Bartonella* prevalence at both sites (*R. satarae*, *M. cf. fernandoni*, *M. cf. famulus*, and *M. cf. terricolor*) and overall community at both sites (**Figure 3A**). Since samples were collected in different years (2018 in Kadamane and 2021 in Kudremukh), we did not attempt any comparison between sites, but treated them separately. Pooled ectoparasite positivity was largely correlated with prevalence in small mammals, albeit slightly lower in every case. Positivity in specific ectoparasite species is provided in **Table S3.**

**Figure 3:**
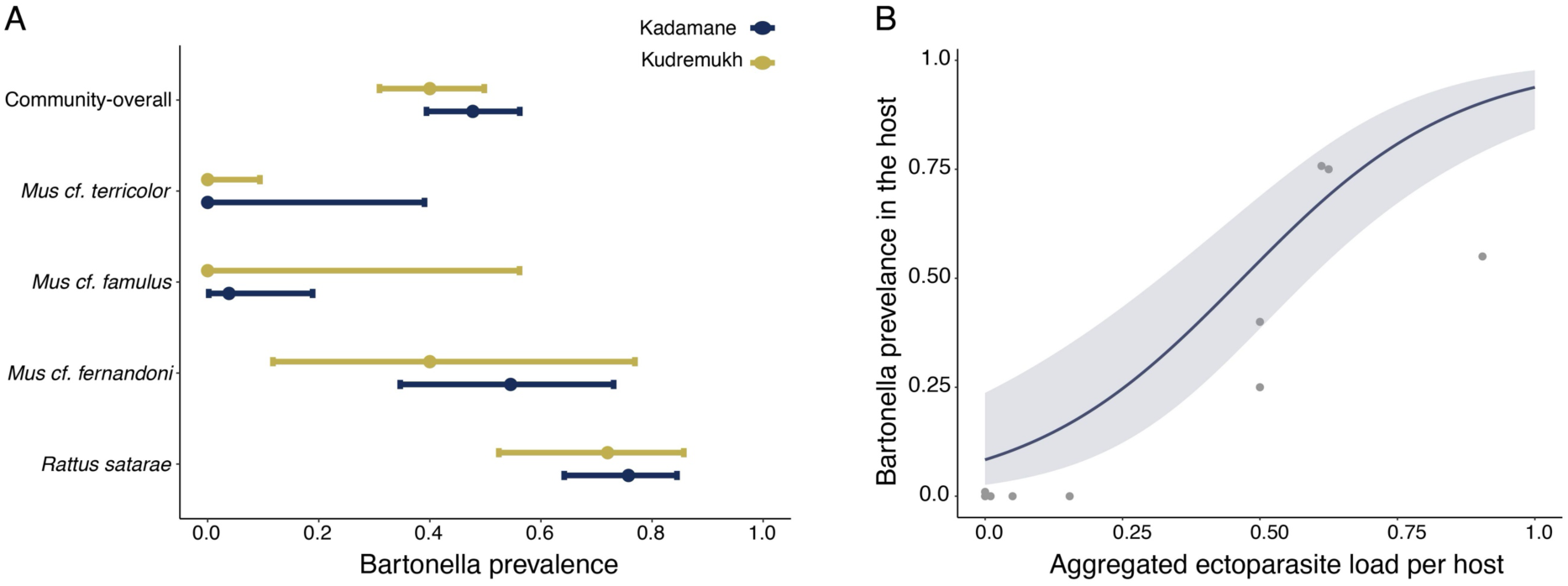
(A) *Bartonella* prevalence in the overall community and the four most abundant small mammals. (B) *Bartonella* prevalence positively correlated with aggregated ectoparasite densities within a small mammal species at each habitat site. The trend line indicates predicted *Bartonella* prevalence as a function of aggregated ectoparasite load. The shaded area represents the 95% confidence interval of the predicted prevalence estimate.

### (e) Factors affecting *Bartonella* prevalence

*Bartonella* prevalence increased with aggregated ectoparasite load (mites, ticks, and fleas) within a species at each habitat site (GLM, p<0.05, **Figure 3B**). We did not find evidence that *Bartonella* prevalence was influenced by host densities and sampling locations (**Table 2**). A second GLM with host species identity and habitat type as additional covariates did not converge; hence we tested these variables separately and found them not significant.

**Table 2:** Results of generalised linear model (GLM) examining drivers of *Bartonella* prevalence. Significant variables are marked with (*)

|  | Estimate | Std. Error |
| --- | --- | --- |
| Intercept | -1.633595 | 1.05266 |
| Host density | -0.07574 | 0.06711 |
| Site | -0.47880 | 0.72288 |
| Aggregated ectoparasite load | 5.10933* | 0.96562 |

### (f) Phylogenetic relationship

Phylogenetic analysis (*rpoB*) revealed eight distinct *Bartonella* lineages (BL1-BL8) from small mammals and ectoparasites. We found that five lineages among them (BL1-BL5) were phylogenetically related to known zoonotic *Bartonella* species (**Figure 4**). We observed three lineages in *R. satarae* (BL1, BL6, and BL7) and two lineages in *M. cf. fernandoni* (BL4 and BL8). BL1 included *Bartonella* sequences from *R. satarae* and ectoparasites (*Laelaps sp 1, Rhipicephalus sp, Ixodes sp, Xenopsylla sp 1, and Xenopsylla sp 5*). Interestingly, sequences from *Laelaps sp 1* and *Xenopsylla sp 5* in this lineage were not collected from *R. satarae* but *M. cf. fernandoni* and *S. niger*, respectively. BL2 consisted of a single sequence collected from *R. rattus*, and closely related to BL1. These two lineages were phylogenetically related to *B. queenslandensis*. BL3 consisted of two sequences, one from *S. niger* and the other from *Laelaps sp 1* collected from *M. cf. fernandoni*. This lineage was related to *B. tribocorum*. BL4 consisted of *Bartonella* sequences from *M. cf. fernandoni*, *M. cf. famulus,* and ectoparasites (*Laelaps sp 1, Rhipicephalus sp, Ixodes sp, Haemaphysalis sp, and Xenopsylla sp 3*). In this lineage, the *Haemaphysalis sp* was recovered from *R. satarae* and not from any of the *Mus* species. BL5 was a single sequence from *F. tristriatus* (reported in Ansil et al. 2021 [27], accession number-MT787671), forming a sister lineage to BL4. We included this sequence in our analysis since it was collected during our study in Kadamane. The remaining lineages (BL6-BL8) were related to non-pathogenic *Bartonella* species. BL6 comprised sequences from *R. satarae* (second lineage from *R. satarae*) and *Ixodes sp* recovered from them. This lineage was related to *B. phoceensis*. BL7 and BL8 were two distinct lineages from *R. satarae* (third lineage from *R. satarae*) and *M. cf. fernandoni* (second lineage from *M. cf. fernandoni*), respectively. Both these lineages were phylogenetically related to *B. thailandensis*.

**Figure 4:**
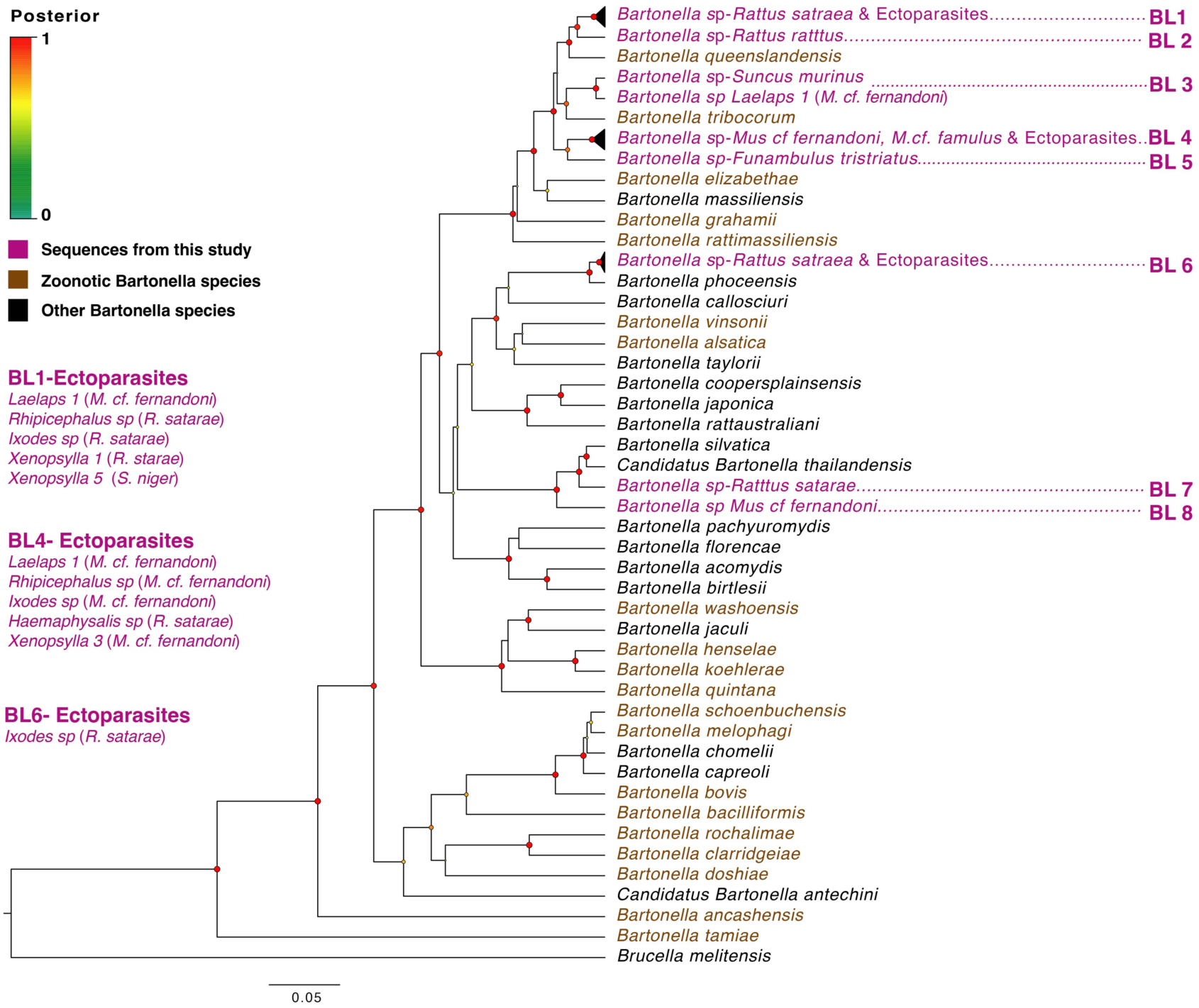
Bayesian phylogenetic inference of *Bartonella* based on 773 bp of *rpoB* sequences. Nodes are coloured based on posterior probability, and monophyletic clades are collapsed for easy visualization. Ectoparasites part of collapsed lineages and their hosts are expanded on the left side.

### (g) Host-*Bartonella* association and historic spillover

The co-phylogeny of small mammals and *Bartonella* revealed host-specific lineages (**Figure 5A**). However, one *Bartonella* lineage from *M. cf. fernandoni* (BL8) was observed to be phylogenetically related to *Bartonella* sequences from *R. satarae* (BL7). Our phylogenetic ancestral trait reconstruction revealed a high probability (0.87) of *R. satarae* being the host state of the common ancestor of these lineages (**Figure 5B**), suggesting a potential past spillover event. Similarly, we also observed a moderately high probability (0.73) of *R. satarae* being the host state for the *Bartonella* sequence from *R. rattus* and the remote common ancestor (0.8) of the *Bartonella* community level phylogeny. We found no evidence for contemporary *Bartonella* spillover or sharing between small mammal hosts.

**Figure 5:**
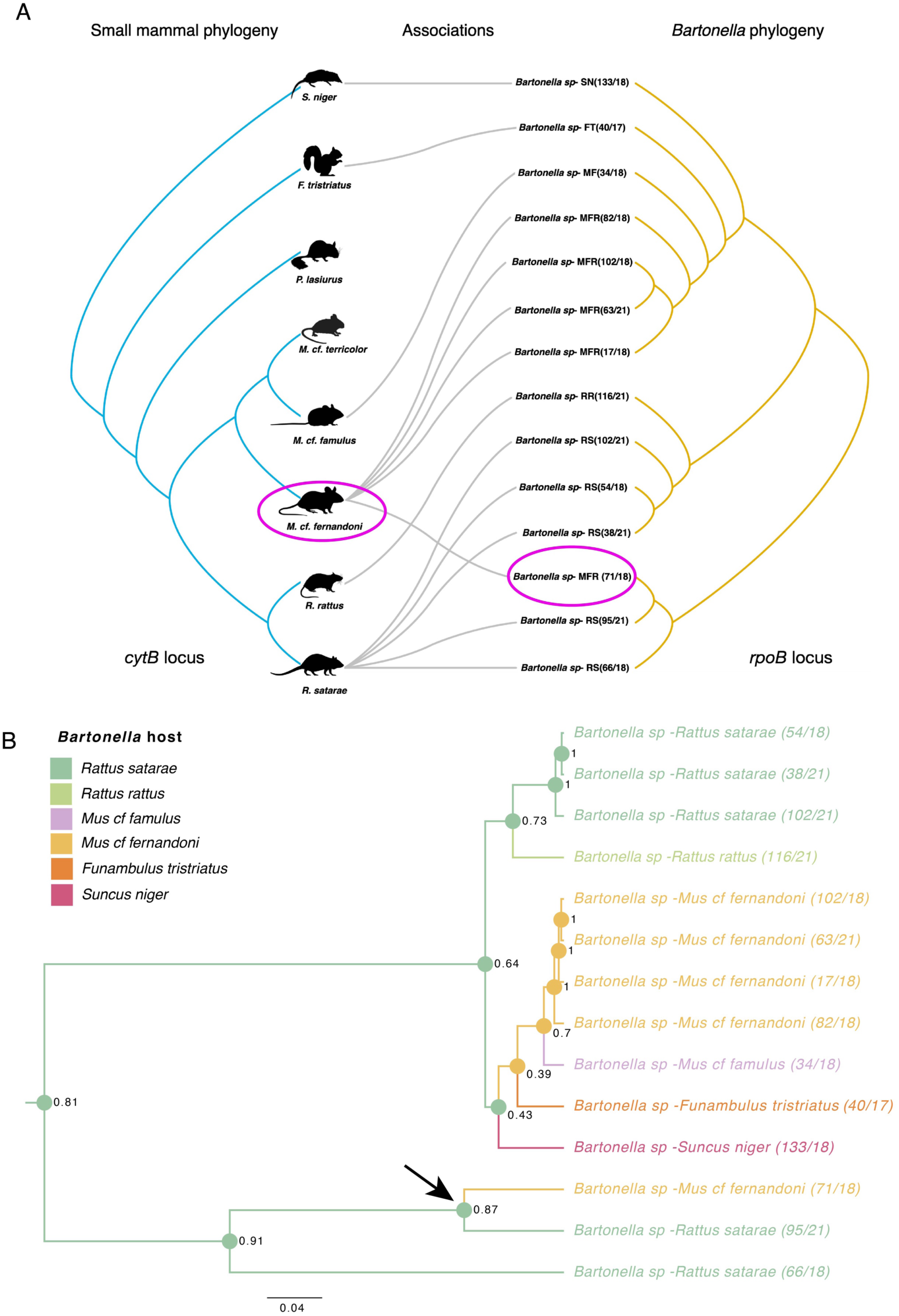
A) Co-phylogeny of small mammals and *Bartonella* association. One lineage of *M. cf. fernandoni* (BL8) was observed to have phylogenetic similarity with one of the *R. satarae* lineage (BL7). B) Bayesian phylogenetic inference showing ancestral host state of different *Bartonella* haplotypes. Each colour represents a specific host, and the number at the common ancestor nodes indicates the probability for the corresponding host state.

## 4. Discussion

We characterized small mammal communities, their ectoparasites, and associations with *Bartonella* in a human-dominated rainforest-plantation landscape in the Western Ghats. We demonstrated that certain small mammal host species occur across habitats, bringing them in contact with specialist hosts with specific *Bartonella* lineages. We also demonstrate that ectoparasites can serve as potential vectors, because *Bartonella* prevalence in the community is influenced by the cumulative ectoparasite load in small mammals. Additionally, we present evidence of a historic spillover event between two major hosts in the community and substantiate the potential for future *Bartonella* sharing between hosts, including other domestic animals and humans, possibly through ectoparasites.

### (a) Some small mammals have the potential to carry *Bartonella* across habitat boundaries

We observed strong species-habitat associations in the focal small mammal community, characterised by exclusive occurrences and high densities, mirroring several ecological communities [54,55]. While species like *R. satarae* (a major *Bartonella* host with highest prevalence) showed high abundance and nearly exclusive occurrence in forested habitats, *R. rattus*, a known synanthropic species, was only detected in built-up areas, reaffirming its commensal nature [56]. Similarly, *M. cf. terricolor* was exclusively recorded in grasslands in both sites, albeit with higher density in Kudremukh. Other *Mus* species, *M. cf. fernandoni* and *M. cf. famulus* were also abundant in grasslands, indicating a generalizable association of *Mus* species with grassland ecosystems. Significantly from a disease ecology perspective, few species like *M. cf. famulus* and *S. niger*, however, were recorded in multiple habitats, including forests, grasslands, and built-up areas. Species like *M. cf. famulus* can also be incidental hosts (as may be the case with the single positive), connecting potential reservoir hosts like *R. satarae* and *M. cf. fernandoni*, with *R. rattus* (another seemingly incidental host with a single positive) and humans. Similarly, *M. cf. fernandoni* was found to occur at low densities in forests in addition to grasslands, bringing them in contact with *R. satarae*, potentially paving the way for the historic spillover (discussed later) from *R. satarae* to *M. cf. fernandoni*.

We did not find any notable difference between small mammal communities within Kudremukh (protected area) and Kadamane (forest-plantation mosaic). One reason for this might be that both landscapes (and much of the Western Ghats) have a long history of human use [25,57], dampening differences that might exist between ‘natural’ and human dominated areas. However, inadequate temporal replicates and delay in sampling Kudremukh diminishes the generality of the pattern observed. Interestingly, all *Mus* species in Kadamane showed variation in annual densities, indicating annual population oscillations [58,59]. We could not fully capture this pattern due to the relatively short duration of this study. However, understanding such variation in host density is crucial as it can be a determinant of zoonotic hazard and spillover [8,60]. This reiterates the importance of detailed ecological studies on host communities to understand spillover dynamics.

Our study suggests that small mammals in this part of the Western Ghats are in urgent need of taxonomic revision. Previous studies from the landscape morphologically identified *M. cf. fernandoni* as *M. musculus* and *M. cf. famulus* and *M. cf terricolor* as *M. booduga* [27]. Phylogenetic analysis and genetic distances suggest that these are distinct lineages related to *M. fernandoni*, *M. famulus*, and *M. terricolor*, respectively. We urge researchers to sample rodents in this landscape and resolve their taxonomy on priority, because pathogen and habitat associations are often linked with species identity as we show in this study.

### (b) *Bartonella* prevalence and spillover may be mediated by ectoparasites

Our results have revealed some broad ectoparasite-host associations that may have implications for *Bartonella* maintenance and transmission. The observed patterns include instances of polyparasitism, host sharing, and the influence of ectoparasitism on *Bartonella* prevalence. In our dataset, ectoparasite diversity and density was centered around medium to large-bodied, and synanthropic species. Specifically, *R. satarae* and *M. cf. fernandoni*, two relatively large-bodied and abundant species with high *Bartonella* prevalence, displayed polyparasitism, often with high densities of specific ectoparasites. This pattern aligns with the positive correlation between body size and ectoparasitism observed in other studies [61]. Potentially, habitat sharing of these species with other small mammals facilitates novel ectoparasite acquisitions and driving overall ectoparasite richness on them [62,63].

Host sharing was another predominant pattern in the ectoparasite community. Several ectoparasites (e.g., *Rhipicephalus sp*, *Xenopsylla sp 1*) showed a wider host range, presumably due to their ability to infect multiple locally available hosts [64]. These results are consistent with previous studies and indicate that host sharing among vectors is more common than previously believed [65]. Such non-specific host-ectoparasite associations can facilitate transmission of several pathogens, making their ecology a crucial part of vector-borne pathogen dynamics [66].

Contrary to our expectations based on host density, our results revealed ectoparasitism (aggregated ectoparasite load per species) was the most important predictor for prevalence. While we did not find the patterns we expected for *Bartonella* prevalence, the correlation between ectoparasitism and *Bartonella* prevalence holds ecological and epidemiological significance. *Bartonella* is a vector-borne pathogen known for its transmission through blood-feeding arthropods [67]. Hence, elevated densities of infected ectoparasites in the environment (and on a specific host) can increase transmission (density-dependant transmission) and impact prevalence and risk of infection [68]. Polyparasitism and lack of host specialization suggests high chances of *Bartonella* transmission between hosts through ectoparasites. In addition, known synanthropic species such as *R. rattus* with multiple generalist fleas (occur in many hosts as demonstrated in this study) pose risks of *Bartonella* transmission to humans and domestic animals [67].

Unfortunately, ecological drivers of aggregated ectoparasite load remain poorly understood. As discussed earlier, longitudinal sampling targeting larger sample sizes from each representative species in combination with ecological models incorporating additional appropriate variables (e.g., host traits, season, etc.) might help explain variance in ectoparasitism. We caution that other variables previously shown to influence pathogen prevalence, like seasonality [9] and environmental stressors [69] were unavailable in our study.

### (c) Non-specific ectoparasite-*Bartonella* associations and evidence of historic spillover suggest future spillover risk

Characterizing *Bartonella* diversity and its association with small mammals and ectoparasites is crucial from a public health perspective, especially since many of the *Bartonella* species are zoonotic and known to associate with small mammals and their ectoparasites [70]. For instance, endemic *Bartonella* variants such as *B. bacilliformis*, responsible for Carrion’s disease, can be fatal and cause serious mortality [71]. Five (of eight) *Bartonella* lineages identified in this study share a close evolutionary relationship with known zoonotic species such as *B. queenslandensis* and *B. tribocorum*. Previous studies have documented the widespread presence and global circulation of these *Bartonella* species, primarily facilitated by small mammals and domestic animals [72]. However, it is important to note that phylogenetic similarity alone does not confirm their ability to cause disease in humans. Additional validation through molecular and *in vitro* experiments is warranted to establish their pathogenicity in humans [73,74].

We observed contrasting patterns in *Bartonella* association with small mammals and ectoparasites. In hosts, *Bartonella* displayed species-specific associations, as we expected based on previous studies [27,75]. Remarkably, *Bartonella* showed non-specific association with ectoparasites (e.g*., Xenopsylla sp* 5 collected from *S. niger* carried *Bartonella* associated with *R. satarae*), leading to a higher overall diversity of haplotypes in specific ectoparasites compared to their corresponding hosts. These complex patterns are not entirely unprecedented, as they have been documented elsewhere [76]. Nevertheless, these non-specific associations between *Bartonella* and ectoparasites have significance for cross-species transmission, given vector competency.

Our phylogenetic ancestral trait reconstruction also highlighted the possibility for cross-species transmission in the community. We observed *R. satarae* as the ancestral host for several *Bartonella* lineages with a medium to high probability, indicating past spillover events involving this species. This was evident between the BL7 and BL8 lineages, recovered from *R. satarae* and *M. cf. fernandoni*, respectively. Interestingly, these two species stood out as the primary *Bartonella* hosts in our dataset, displaying high prevalence, high haplotype diversity, and co-occurrence in forested habitats. Such pathogen movement between host species could provide an opportunity to evolve new virulent variants through recombination [4], similar to those observed in other pathogen systems [6]. Although we did not find instances of contemporary spillover, our results suggest past spillover events, potentially through ectoparasites, emphasizing the likelihood of similar events. The communities we studied include many small mammals and their ectoparasites that overlap between habitats, increasing the likelihood of such inter-species transmissions [77]. In addition, the movement of generalist hosts across multiple habitats carrying generalist ectoparasites with non-specific *Bartonella* associations is likely to further drive pathogen spillover in the landscape [78].

## 5. Conclusion

This study has uncovered several significant insights and nuances about the intricate relationship between small mammal hosts, their ectoparasites, and bacterial agents within a tropical mixed-use landscape. While our model system and investigation were specific to *Bartonella*, the results and inferences extend to a broader spectrum of vector-borne bacterial pathogens in multi-host communities. Our research highlights the crucial role of ectoparasitism in driving *Bartonella* prevalence within these small mammal communities where some hosts move across habitat boundaries. Our results also highlight the non-specific association of ectoparasites with *Bartonella* diversity compared to their hosts, underscoring the potential for ectoparasite-driven cross-species transmissions. Using evolutionary analyses, we further demonstrate the role of historical spillover events in shaping the current *Bartonella* distribution in the community and highlight the future prospects of such spillover events. Collectively, our findings underscore the need for comprehensive ecological studies to advance our understanding of the transmission dynamics of vector-borne bacterial pathogens within multi-host communities and their zoonotic implications, particularly in rapidly changing landscapes.

## Supporting information

Supplementary results

## Ethics statement

This study and associated protocols were approved by National Centre for Biological Science Institutional Animal Ethics Committee (NCBS-IAEC-2016/10-[M], NCBS-IAE-2020/ 02[N]) and Institutional Biosafety Committee (TFR:NCBS:23_IBSC/2017).

## Data accessibility

The manuscript and Supplementary materials present small mammal capture data and *Bartonella* prevalence data from small mammals and ectoparasites. *Bartonella* sequence data generated during this study are deposited in GenBank under the accession numbers OR574780-OR574826.

## Author contributions

ABR: conceptualization, methodology, investigation, data curation, formal analysis, visualization, project administration, writing -original draft, writing-review and editing; AV: methodology, data curation, formal analysis, visualization, writing-review and editing; VR: methodology, investigation, writing-review and editing; HMY: methodology, investigation, writing-review and editing; AS: investigation, writing-review and editing; UR: conceptualization, funding acquisition, project administration, supervision, writing-review and editing.

## Acknowledgement

This study was primarily supported by the Department of Atomic Energy, Government of India (Project Identification RTI 4006) awarded to UR. We are grateful to Mr. K.M. Cariappa of Kadamane for facilitating the field study; Karnataka Forests and Wildlife Department and Kudremukh Wildlife Division for granting permission for small mammal sampling; Kudremukh Iron Ore Company for providing accommodation in Kudremukh. We thank S. Yashas, Vikrant Jathar, S. Vijayakumar, K. Kamaraj, Tamil Selvan, Subramanian, and C. Karnan for their help in field sampling. We also acknowledge the support of NCBS Sanger sequencing facility and museum and field Stations facility. ABR was supported by the SPM fellowship by CSIR-HRDG, Government of India. ABR also acknowledges the support from the Tata Institute for Genetics and Society during the manuscript preparation. AS was supported by fellowship from the Trivedi School of Biosciences, Ashoka University.

