## Supplementary results for "Host-pathogen-vector continuum in a changing landscape: Drivers of *Bartonella* prevalence and evidence of historic spillover in a multi-host community"

### a) Small mammal density

We were able to estimate densities for four (of eleven) common small mammal species (*Rattus satarae*, *Mus cf. fernandoni*, *Mus cf. famulus* and *Mus cf. terricolor* ). These densities are shown in Figure S2. *R. satarae* was only captured in forests (except on one occasion in Kudremukh grassland) and showed similar density (5 individuals/hectare) in both Kadamane and Kudremukh (**Figure S2A**). Given similar vegetation structure, we included rested plantations as forests for all our analyses. In 2018, 13 individuals *R. satarae* (per hectare) were captured, but there were no recaptures (see methods) to estimate density. *M. cf. fernandoni* showed high density (13 individuals/hectare) in Kadamane grassland in 2016 (**Figure S2B**). This species was captured in small numbers in other years, making the density estimation challenging. *M. cf. famulus* captured in all land-use types (**Table S1**), however showed high density (16 individuals/hectare) in Kadamane grasslands in 2016 (**Figure S2C**). *M. cf. terricolor* was captured only in grasslands and showed highest density (80 individuals/hectare) in Kudremukh (**Figure S2D**). All the three *Mus species* showed signs of annual population fluctuations in grasslands.

**b) Ectoparasite density**

A high density of *Laelaps sp 1* was observed in *M. cf. fernandoni* from forests (27.5, n=2) and grasslands (7.2, n=27; **Figure S3A**). *Laelaps sp 2* was only detected from *P. lasiurus* (3.5, n=2), an arboreal small mammal endemic to the Western Ghats. Mites were very rare/absent (0.08, n=91) on *R. satarae*. Ticks were only detected on small mammals captured from forests and grasslands. In forests, *R. satarae* showed a high density of *Ixodes sp* (1.2, n=91), followed by *Haemaphysalis sp* (0.5, n=91) and *Rhipicephalus sp* (0.3, n=91; **Figure S3B**). In grasslands, *M. cf. fernandoni* also carried all three ticks; *Rhipicephalus sp* (0.5, n=27)*, Ixodes sp* (0.1, n=27) and *Haemaphysalis sp* (0.1, n=27) while *M. cf. terricolor* carried only *Rhipicephalus sp* (0.2, n=45) and *Haemaphysalis sp* (0.2, n=45). On one occasion, when *R. satarae* and *Suncus niger* was caught in the grassland, a large number of *Ixodes* ticks (27 and 17, respectively) were attached to them. These unusual numbers are manually represented in Figure S2B to maintain the spread along the Y-axis. Surprisingly, no mites and ticks were detected from small mammals from built-up areas.

Among the five species of *Xenopsylla* species detected, two were seen in forests; *Xenopsylla sp3* (1.5, n=3) and *Xenopsylla sp5* (0.7, n=2), associated with *M. cf. fernandoni* and *S. niger*, respectively (**Figure S3C**). In the grassland, *Xenopsylla sp1* was detected from *R. satarae* (1, n=1) and *Xenopsylla sp3* from *M. cf. fernandoni* (0.1, n=27). In the built-up area, all fleas were recorded except *Xenopsylla sp5*. *R. rattus* carried *Xenopsylla sp1* (0.1, n=14), *Xenopsylla sp2* (0.1, n=14) and *Xenopsylla sp4* (0.2, n=14). *F. tristriatus* carried *Xenopsylla sp1* (1, n=1) and *Xenopsylla sp3* (3, n=1; which is manually marked in the figure to highlight the variation between smaller density values) while *S. niger* carried *Xenopsylla sp3* (0.2, n=5) alone.

**c) *Bartonella* positivity in ectoparasites**

Ectoparasite pools showed 32.4% (n=102) and 15.6% (n=45) *Bartonella* positivity in Kadamane and Kudremukh, respectively **(Table 1)**. Seven of the ten ectoparasite morphotypes showed *Bartonella* positivity; *Laelaps sp1*, *Rhipicephalus sp*, *Ixodes sp*, *Haemaphysalis sp*, *Xenopsylla sp 1*, *Xenopsylla sp 3,* and *Xenopsylla sp 5* **(Table S3).** We observed high positivity in *Ixodes sp* (58.3, n=36) followed by *Rhipicephalus sp* (25%, n=25) and *Laelaps sp1* (22.2%, n=27). Other species, such as *Xenopsylla sp 1* (15%, n=20), *Xenopsylla sp 3* (16.7%, n=6), and *Haemaphysalis sp* (3.8%, n=26) had relatively lower positivity. Morphotypes such as *Xenopsylla sp 2*, *Xenopsylla sp 4*, *Xenopsylla sp 5*, and *Laelaps sp 2* had smaller sample sizes among the pools, hence true positivity is uncertain.
