## Supplementary tables for "Host-pathogen-vector continuum in a changing landscape: Drivers of *Bartonella* prevalence and evidence of historic spillover in a multi-host community"

**Table S1:** Small mammals captured in different land-use types across different years. Year 2016, 2017, and 2018 represent samples from Kadamane and 2021 represents samples from Kudremukh National Park.

| **Species** | **Forest** | | | | **Grassland** | | | | **Tea plantation** | | | **Built-up** | | | |
|---|---|---|---|---|---|---|---|---|---|---|---|---|---|---|---|
| | **2016** | **2017** | **2018** | **2021** | **2016** | **2017** | **2018** | **2021** | **2016** | **2017** | **2018** | **2016** | **2017** | **2018** | **2021** |
| *Rattus satarae* | 21 | 13 | 67 | 24 | - | - | - | 1 | - | - | - | - | - | - | - |
| *Rattus rattus* | - | - | - | - | - | - | - | - | - | - | - | 4 | 2 | 10 | 4 |
| *Mus cf. fernandoni* | 15 | 13 | 2 | - | 22 | 1 | 21 | 6 | - | - | - | - | - | - | - |
| *Mus cf. famulus* | 1 | 2 | 2 | - | 27 | 2 | 20 | 3 | 3 | - | - | - | 1 | 5 | - |
| *Mus cf. terricolor* | - | - | - | - | 40 | 5 | 6 | 39 | - | - | - | - | - | - | - |
| *Golunda ellioti* | - | - | - | - | 1 | 2 | - | - | - | - | - | - | - | - | - |
| *Vandeleuria nilagirica* | 1 | - | - | - | - | - | - | - | - | - | - | - | - | - | - |
| *Platacanthomys lasiurus* | - | 2 | 1 | 1 | - | - | - | - | - | - | - | - | - | - | - |
| *Funambulus tristriatus* | - | 1 | - | - | - | - | - | - | - | - | - | - | - | - | 1 |
| *Suncus niger* | - | - | - | 3 | - | - | - | 1 | - | - | - | - | 1 | 2 | 3 |
| *Crocidura horsfieldii* | - | - | - | - | 3 | 1 | - | - | - | - | - | - | - | - | - |
| **Total** | **38** | **31** | **72** | **28** | **93** | **11** | **47** | **50** | **3** | **-** | **-** | **4** | **4** | **17** | **8** |

**Table S2:** Ectoparasite occurrence in small mammal species. The number indicates number of occasions a specific morphotype (species) was observed in a host species. The percentage indicate ectoparasite load on a specific species (equivalent to prevalence).

| | **Ectoparasites occurrence in small mammals (%)** | | | | | | | | |
|---|---|---|---|---|---|---|---|---|---|
| | *Rattus satarae n=92* | *Rattus rattus n=14* | *Mus cf. fernandoni n=29* | *Mus cf. famulus n=30* | *Mus cf. terricolor n=45* | *Platacanthomys. lasiurus n=2* | *Funambulus tristriatus n=1* | *Suncus niger n=9* | Total *n=222* |
| *Laelaps sp 1* | 3 (3.3) | - | 24 (82.8) | - | - | - | - | - | 27 (12.2) |
| *Laelaps sp 2* | - | - | - | - | - | 1 (50) | - | - | 1 (0.5) |
| *Rhipicephalus sp* | 18 (19.6) | - | 4 (13.8) | - | 5 (11.1) | - | - | 1 (11.1) | 28 (12.6) |
| *Ixodes sp* | 34 (37) | - | 2 (6.9) | - | - | - | - | - | 36 (16.2) |
| *Haemaphysalis sp* | 22 (23.9) | - | 2 (6.9) | - | 2 (4.4) | - | - | - | 26 (11.7) |
| *Xenopsylla sp 1* | 18 (19.6) | 1 (7.1) | - | - | - | - | 1 (100) | - | 20 (9) |
| *Xenopsylla sp 2* | - | 1 (7.1) | - | - | - | - | - | - | 1 (0.5) |
| *Xenopsylla sp 3* | - | - | 3 (10.3) | 1 (3.3) | - | - | 1 (100) | 1 (11.1) | 6 (2.7) |
| *Xenopsylla sp 4* | - | 1 (7.1) | - | - | - | - | - | - | 1 (0.5) |
| *Xenopsylla sp 5* | - | - | - | - | - | - | - | 1 (11.1) | 1 (0.5) |

**Table S3:** *Bartonella* positivity in various ectoparasites (pooled) recovered from small mammal species. Since ectoparasites collected were pooled for screening, positivity reported here is not true prevalence in the ectoparasite community.

| | ***Bartonella* positivity in pooled ectoparasites; number of positives/number of pools tested (%)** | | | | | | | | Total |
|---|---|---|---|---|---|---|---|---|---|
| | *Rattus satarae* | *Rattus rattus* | *Mus cf. fernandoni* | *Mus cf. famulus* | *Mus cf. terricolor* | *Platacanthomys lasiurus* | *Funambulus tristriatus* | *Suncus niger* | |
| *Laelaps sp 1* | 0/3 (0) | - | 6/24 (25) | - | - | - | - | - | 6/27 (22.2) |
| *Laelaps sp 2* | - | - | - | - | - | 0/1 (0) | - | - | 0/1 (0) |
| *Rhipicephalus sp* | 6/18 (33.3) | - | 1/4 (25) | - | 0/5 (0) | - | - | 0/1 (0) | 7/28 (25) |
| *Ixodes sp* | 20/34 (58.8) | - | 1/2 (50) | - | - | - | - | - | 21/36 (58.3) |
| *Haemaphysalis sp* | 1/22 (4.5) | - | 0/2 (0) | - | 0/2 (0) | - | - | - | 1/26 (3.8) |
| *Xenopsylla sp 1* | 3/18 (16.7) | 0/1 (0) | - | - | - | - | 0/1 (0) | - | 3/20 (15) |
| *Xenopsylla sp 2* | - | 0/1 (0) | - | - | - | - | - | - | 0/1 (0) |
| *Xenopsylla sp 3* | - | - | 1/3 (33.3) | 0/1 (0) | - | - | 0/1 (0) | 0/1 (0) | 1/6 (16.7) |
| *Xenopsylla sp 4* | - | 0/1 (0) | - | - | - | - | - | - | 0/1 (0) |
| *Xenopsylla sp 5* | - | - | - | - | - | - | - | 1/1 (100) | 1/1 (100) |
