## Supplementary figures for "Host-pathogen-vector continuum in a changing landscape: Drivers of *Bartonella* prevalence and evidence of historic spillover in a multi-host community"

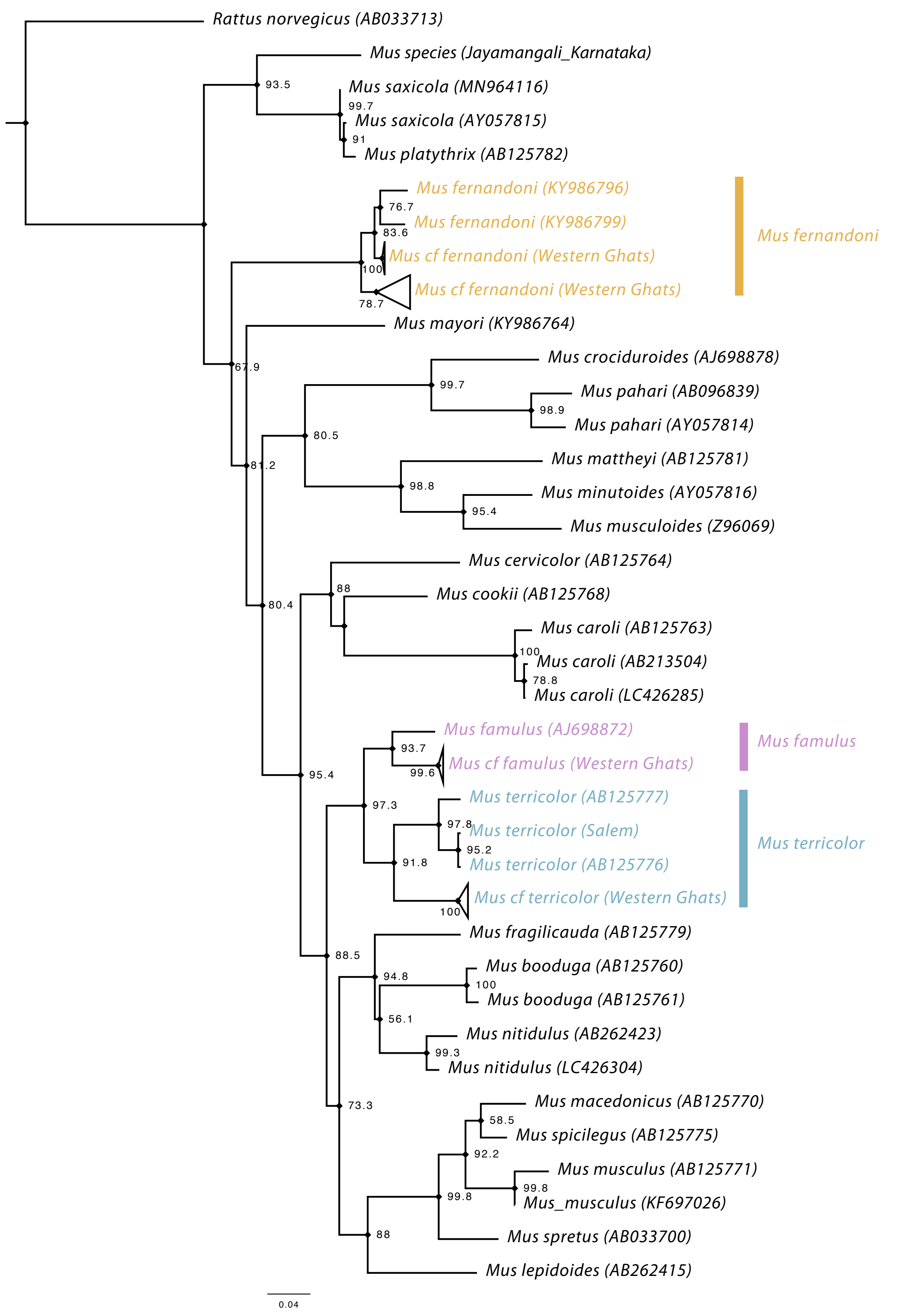


**Figure S1**: Phylogenetic inference of *Mus* species. This phylogenetic tree based on 870 bp of *cytochrome B* gene was used to identify cryptic *Mus species* from the study area (Western Ghats).

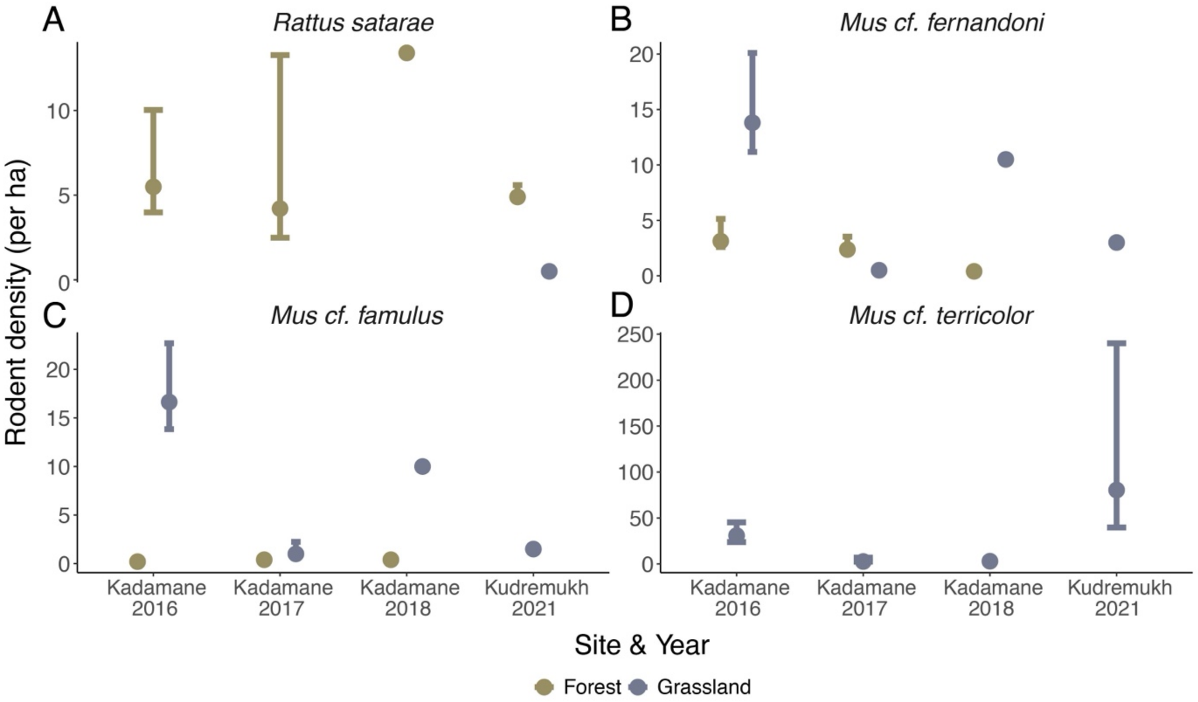


**Figure S2:** Estimated density of the four most common small mammals in the community. Points without error bars indicate number of individuals detected per hectare (due to low sample size or lack of recaptures). In 2018, individuals were sacrificed for organ samples, hence no recaptures for density estimates.

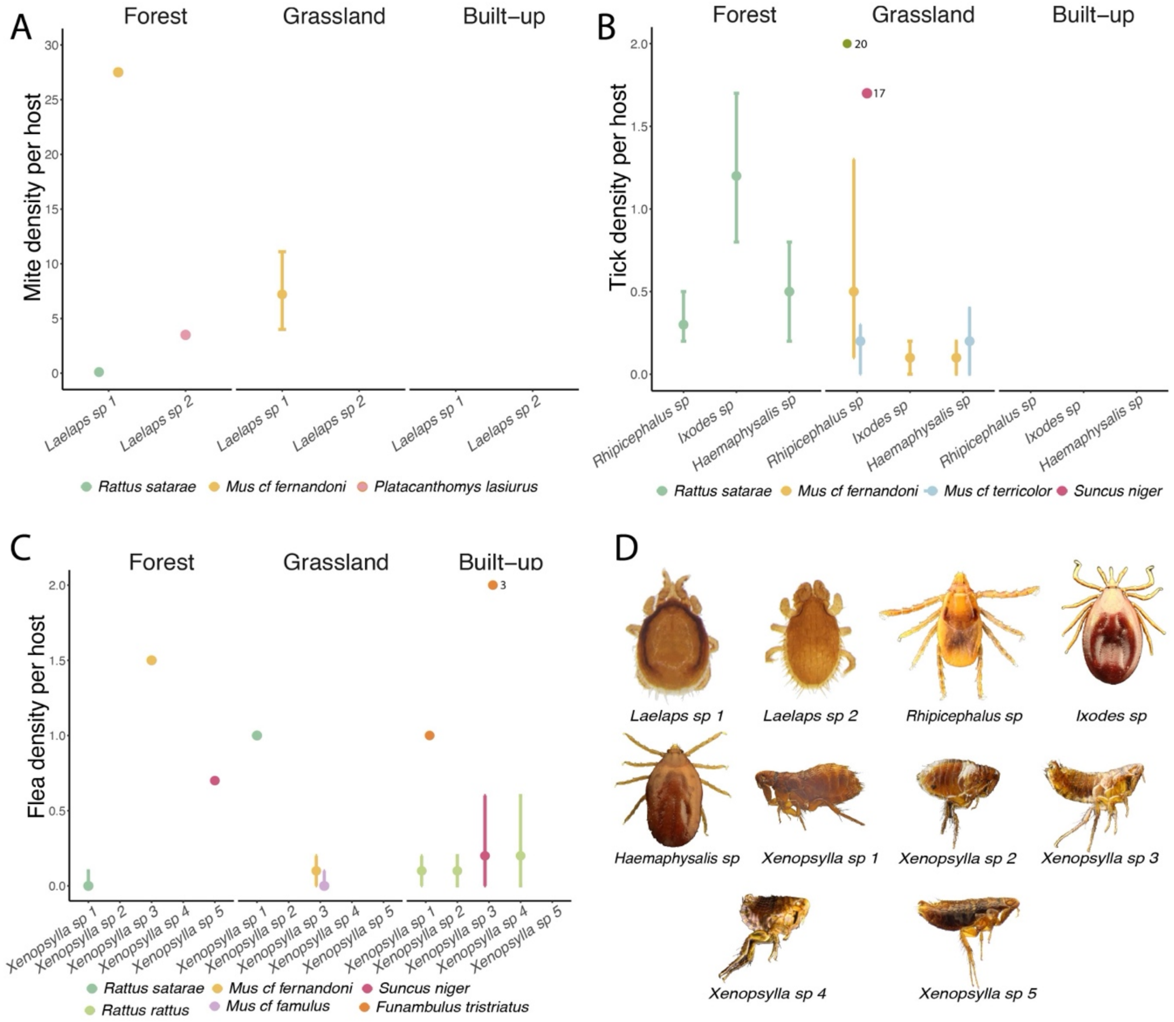


**Figure S3:** Estimated densities of various ectoparasites collected from small mammals. A, B and C show densities of various species of mites, ticks and fleas, respectively. The numbers next to the points in B and C are unusual ectoparasites counts observed manually labelled to preserve Y-axis distribution. D shows microscopic images (not to scale) of different ectoparasites collected from small mammals in the study area.
